# IVIM parameters have good scan-rescan reproducibility when evidential motion contaminated and poorly fitted image data are removed

**DOI:** 10.1101/179440

**Authors:** Olivier Chevallier, Nan Zhou, Jian He, Romaric Loffroy, Yi-Xiáng J. Wang

## Abstract

**Background:** Intravoxel Incoherent Motion (IVIM) diffusion MRI is a promising technique for liver pathology evaluation, but this technique’s scan-rescan reproducibility has been reported to be unsatisfactory.

**Objective:** To understand whether IVIM MRI parameters for liver parenchyma can be good after removal of motion contaminated and/or poorly fitted image data.

**Material and Methods:** Eighteen healthy volunteers had liver scanned twice at the same session to assess scan-rescan repeatability, and again in another session after an average interval of 13 days to assess reproducibility. Diffusion weighted image were acquired with a 3T scanner using respiratory-triggered echo-planar sequence and 16 *b*-values (0 to 800 s/mm2). Measurement was performed on the right liver with segmented-unconstrained least square fitting. Image series with evidential anatomical mismatch, apparent artifacts, and poorly fitted signal intensity vs. *b*-value curve were excluded. A minimum of three slices was deemed necessary for IVIM parameter estimation of a liver.

**Results:** With total 54 examinations, 6 scans did not satisfy inclusion criteria, leading to a success rate of 89%; and 14 volunteers were finally included. With each scan a mean of 5.3 slices (range: 3-10 slices) were utilized for analysis. Using threshold *b*-value=80s/mm2, the coefficient of variation and within-subject coefficient of variation for repeatability and reproducibility were: 2.86% and 4.24% for Dslow, 3.81% and 4.24%, for PF, 18.16% and 24.88% for Dfast; and those for reproducibility were 2.48% and 3.24% for Dslow; 4.91% and 5.38% for PF; 21.18% and 30.89% for Dfast.

**Conclusion:** IVIM parameter scan-rescan reproducibility can be potentially good.

## Introduction

Since the initial study of Yamada et al (1), there have been greater interests to explore Intravoxel Incoherent Motion (IVIM) technique to evaluate diffused liver diseases such as liver fibrosis and nonalcoholic fatty liver disease; to characterize liver tumor; and to evaluate treatment response (2, 3). A prerequisite to translating IVIM imaging into clinical applications is accurate measurement of IVIM parameters and acceptable reproducibility. Nevertheless, accurate liver IVIM quantification is challenging, partially due to the limited sampling and low signal-to-noise ratio (SNR) for fast diffusion data acquisition. A major obstacle for clinical application of IVIM technique for abdominal organs is its unsatisfactory scan-rescan reproducibility (2, 3). For example, in a short-term reproducibility study, Andreou et al (4) reported the 95% confidence intervals of percentage difference between paired measurements of liver parenchyma was (−24.3, 25.1) for PF (*f*), (−5.12, 8.09) for Dslow (*D*), and (−31.2, 59.1) for Dfast (*D*);* and the absolute limit was (0.140, 0.232) for PF, (0.951, 1.08) for Dslow, and (35.7, 82.5) for Dfast.

IVIM diffusing imaging typically involves long data acquisition time (~5 min) with images acquired at a series of *b*-values. It is usually acquired with respiratory gating, however, even respiratory gating is associated with substantial residual respiration induced motion (2). Respiratory motion can cause inter-*b*-value motion and intra-*b*-value motion. Inter-*b*-value motion causes mis-match of anatomical structures on images of difference *b*-values, and intra-*b*-value motion where motion occurs during the data acquisition for the slice causes image artifact/image distortion. Diffusion MRI is also influenced by artifacts related to magnet/sequence imperfections, such as B0 inhomogeneity resulting from susceptibility variations; geometric distortions from residual motion probing gradients-induced eddy currents (5). In this study, we introduce a manual ‘image data cleaning’ process, with the aim to mitigate artifacts associated with respiration motion as well as with magnet/sequence imperfections. We hypothesize that if IVIM’s scan-rescan reproducibility will be satisfactory after ‘image data cleaning’, then with further technical improvement such as acquisition of more *b*-values for curve fitting, advanced methods for motion correction, statistical remove of ill fitted pixels, or accelerated data acquisition with single-breath hold, liver IVIM will eventually have clinical applicability.

## Material and Methods

This study was conducted with the approval of the institutional ethics committee and informed consent was obtained. Eighteen healthy volunteers underwent IVIM diffusion imaging with a 3T magnet and a 32 channels dStream Torso coil (Ingenia, Philips Healthcare, Best, The Netherlands). The IVIM diffusion imaging was based on a single-shot spin-echo-type echo-planar imaging sequence, with 16 b-values of 0, 3, 10, 25, 30, 40, 45, 50, 80, 200, 300, 400, 500, 600, 700 and 800 s/mm2, NSA of 2 for b=700 s/mm2 and b=800 s/mm2, and NSA=1 for other *b*-values. Spectral presaturation with inversion recovery technique was used for fat suppression. Respiratory triggering was performed using an air-filled pressure sensor fixed on the upper abdomen, resulting in an average TR of 2149 ms. Other parameters included TE=55ms, slice thickness=6mm, matrix=100×116, field-of-view (FOV)=360×300 mm, EPI factor= 29, a sensitivity-encoding (SENSE) factor=4, number-of-slices =26. The scan subjects were trained so that they maintained shallow regular breathing during image acquisition. The average IVIM scan duration was 6 min. All volunteers were scanned twice in the same session to assess scan–rescan repeatability (scans 1.1 and scans 1.2), and additionally once again in another session (scan 2) with an interval of 5–21 days (mean 13 days) to assess scan–rescan reproducibility.

The IVIM signal attenuation was modeled according to Eq 1 (6)

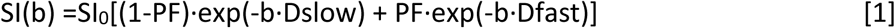

With the segment-unconstrained approach, the estimation of Dslow was obtained by a least-squares linear fitting of the logarithmized image intensity at *b* values greater than threshold, which was chosen of 50, 80 or 200 s/mm^2^, to a linear equation (7). The fitted curve was then extrapolated to obtain an intercept at *b-*value=0 s/mm^2^. The ratio between this intercept and SI^0^ gave an estimate of PF. Finally, the obtained Dslow and PF were substituted into Eq. 1 and were nonlinear least squares fitted against all *b* values to estimate Dfast using the Trust-Region based algorithm.

All curve-fitting algorithms were implemented in a custom program developed on MatLab (Mathworks, Natick, MA, USA). For ROI analysis, the IVIM parameters were calculated based on the mean signal intensity of the whole ROI, which has been shown to offer better estimation than pixel-wise fitting when the signal-to-noise ratio (SNR) of the diffusion weighted images is low (8, 9). The same as some other reports (10-18), only the right lobe of the liver was measured in the current study (Fig. 1).

**Fig. 1.**
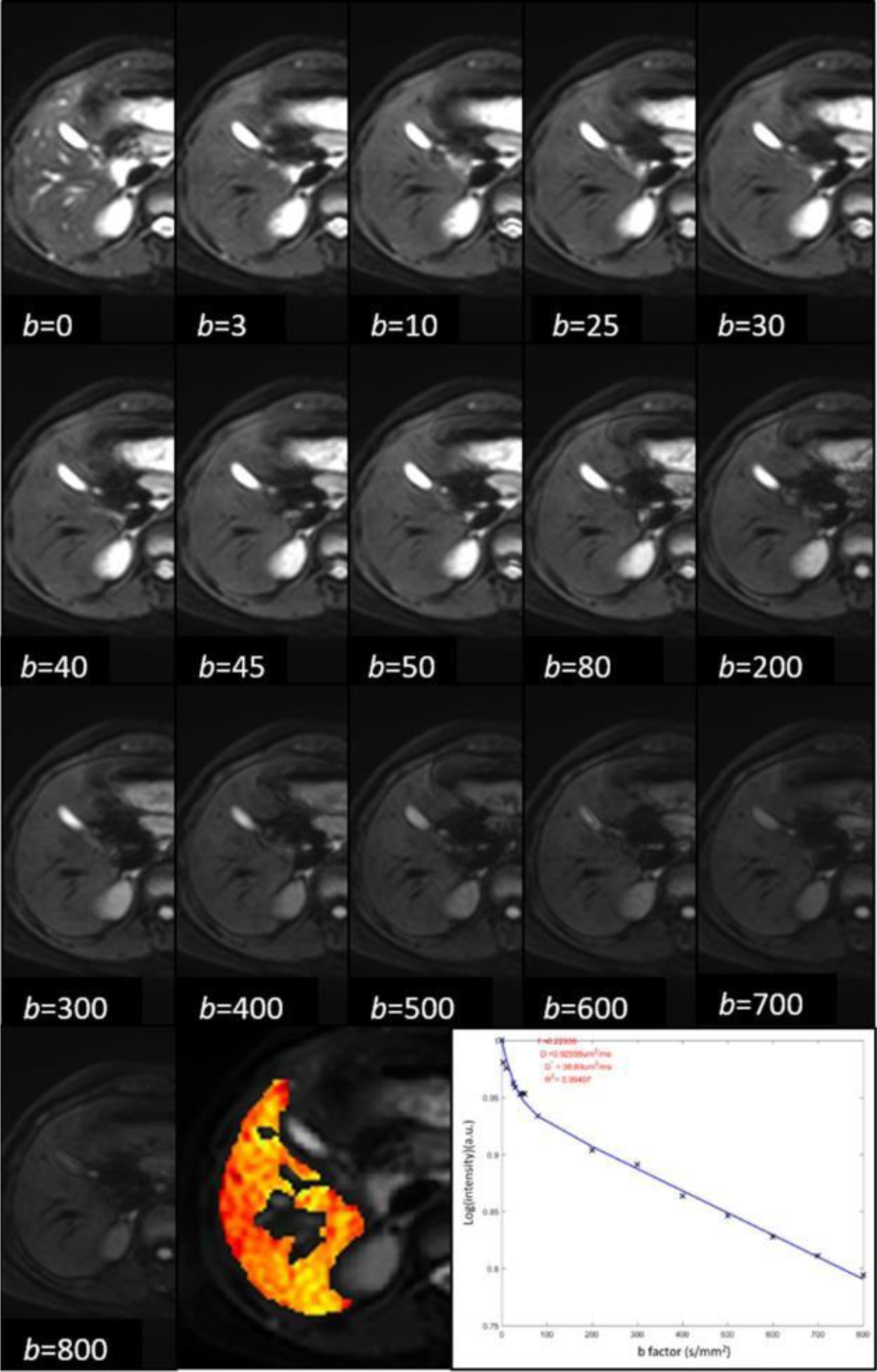
A IVIM image series of ‘good quality’, No evidential motion or artifacts could be seen in more than two images. ROI was drawn to cover as much as possible of liver parenchyma of right lobe, while avoiding signal vessels which are noticeable in the central parts of parenchyma and also in the area close to the gallbladder in this slice. The signal intensity and *b*-value curve has a R^2^>0.96 for ROI-based fitting. The signal decay is consistent with *b*-value changes, and no obvious outlier value is noted.

A manual procedure was taken to ‘clean the image data’ for each examination. Firstly, slices which covered only the lowest part of segment V-VI (usually slices below the gallbladder) or the hepatic dome, near the visceral near or the diaphragmatic surfaces, were discarded. Then, each scan’s image series were graded as ‘good quality’, ‘fair quality’, or ‘insufficient quality’. Motion induced imaging data degrading was visually assessed between consecutives images at different *b*-values for each slice, noting the location of the following anatomic structures: kidneys, gall bladder, spleen, hepatic edges, main hepatic vessels (main portal vein, portal veins until second order, main hepatic veins). If no motion or artifact was noted, the slice series was graded ‘good quality’. Image series of ‘insufficient quality’ were mainly due to motion leading to liver displacement between images of different *b*-values (inter-*b*-value motion), and sometimes apparent artifacts in the hepatic parenchyma which could be due to intra-*b*-value motion (Fig 2). Slices presented only slight displacement or inconspicuous artifact were graded ‘fair quality’. Image series of ‘good quality’ and ‘fair quality’ were included for the second step data cleaning. Image series which generated poorly IVIM diffusion fitted curve were then excluded. Firstly, slices which presented parameters results with a coefficient of determination R^2^ value lower than 0.95 for ROI-wise fitting were excluded (19). Then, the plots of signal intensity vs. *b*-values were individually evaluated. Slices which demonstrated evidential outliers with MRI signal vs. *b*-value relation and could not be properly fitted were discarded. In addition, for an IVIM image series to be valid, we required that at least three slices for each liver examination can be included for final analysis after data-cleaning. The means of all included slices’ measurements were then regarded the value of the examination.

**Fig. 2.**
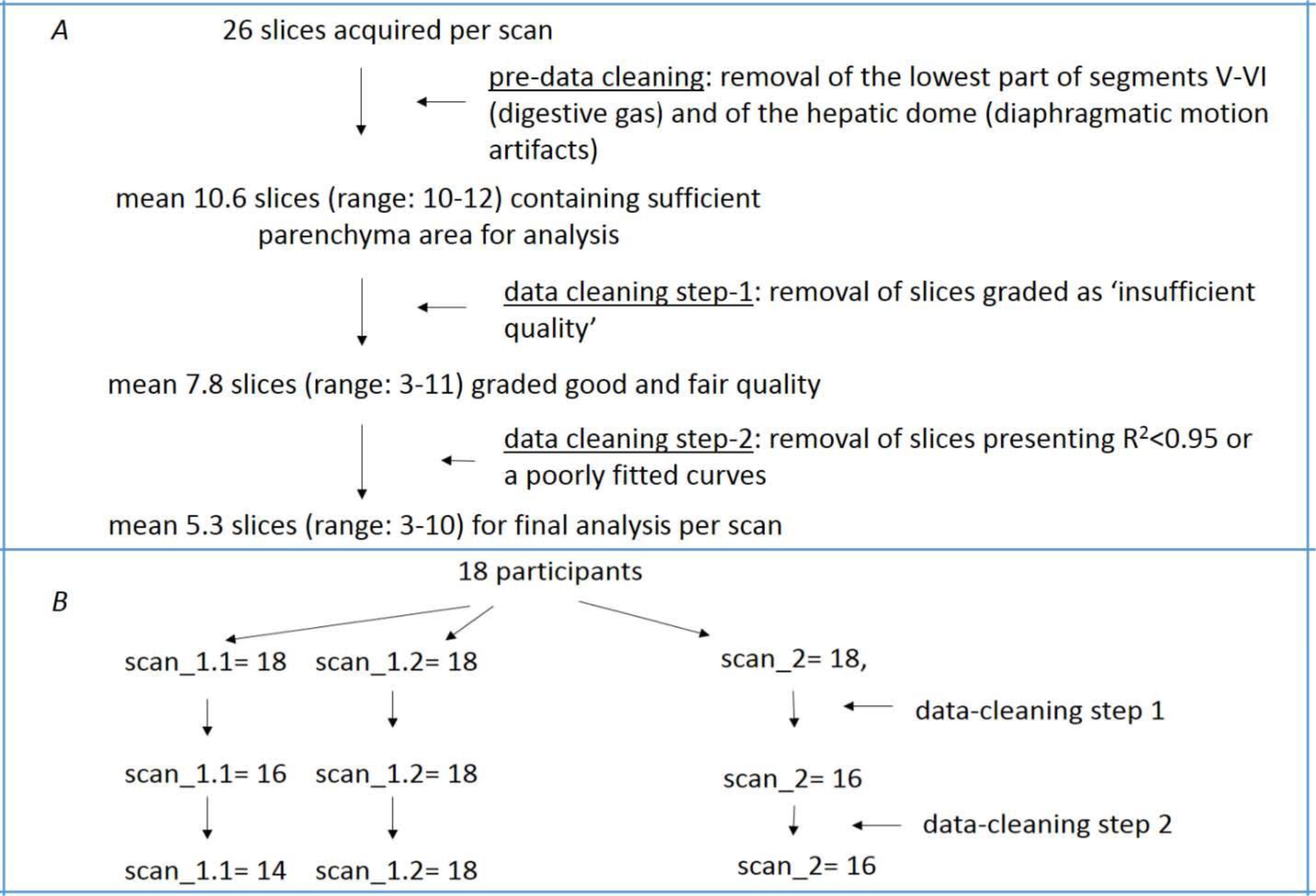
A: image-data cleaning process flow diagram. B: number of volunteer subjected included for analysis at each steps.

**Fig. 3.**
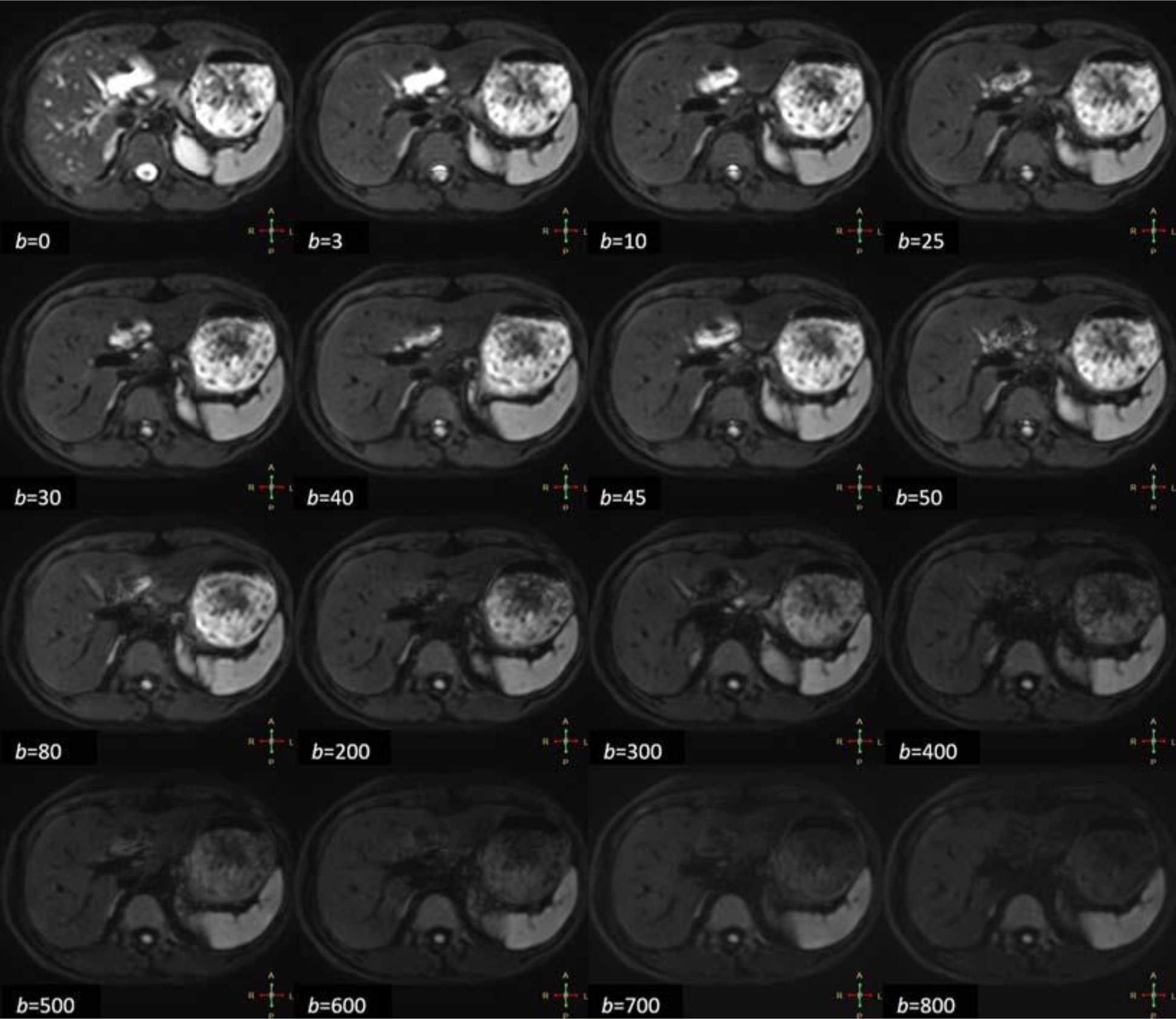
An example of excluded image series due to inter-image motion. Left kidney size varies among different images, and the posterior branch of the right portal branch vein can be seen in some images but not in other images.

**Fig. 4.**
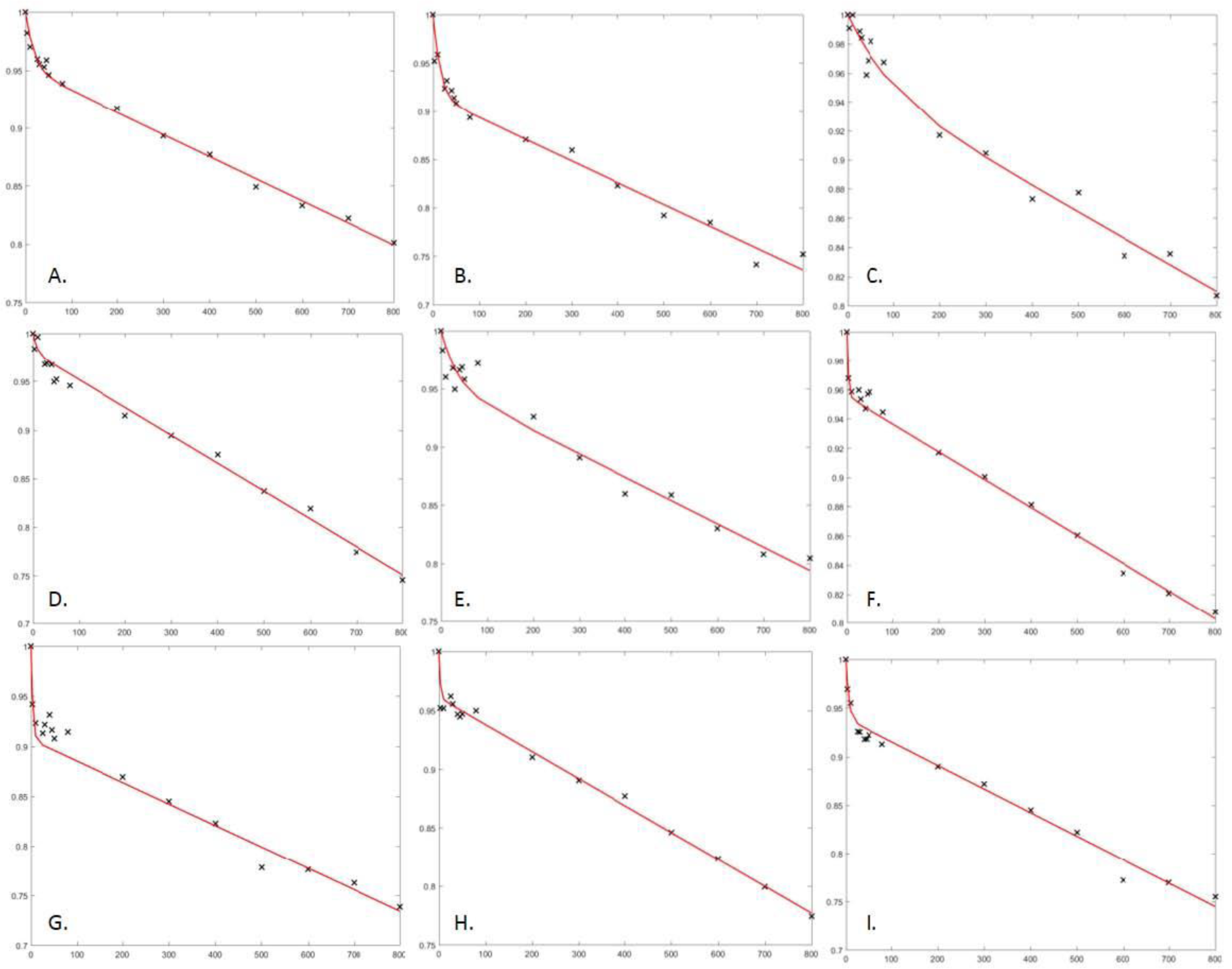
Examples of well fitted (A), acceptable (B) and unacceptable (C-I) signal intensity and *b*-value curves. Plots C, E, F, G, H show MRI signal increases and decreases erratically between *b*=0 s/mm2 and *b*=80 s/mm2. For plots D, F, I, H, at least 3 consecutive data points are outliers under or above the fitted curve between b=0 s/mm2 and b=80 s/mm2. Plots F, G, H present sharp signal drop at *b*-value=0 and b=3 s/mm2, and therefore unreasonably high Dfast values.

Statistical analysis was performed using MedCalc Statistical Software (version 17.6, MedCalc Software bvba, Ostend, Belgium). Intra-scan repeatability between scan 1.1 and scan 1.2, and inter-scan reproducibility between scan 1.1 and scan 2 of PF, Dslow and Dfast were assessed by coefficient of variation (CoV), the within-subject coefficient of variation (wCoV), and Bland-Altman mean difference and 95% limits of agreements (BA-LA). wCoV is defined by Eq 2 and Eq 3

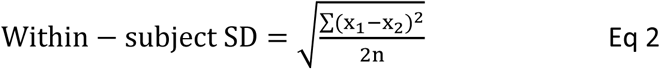

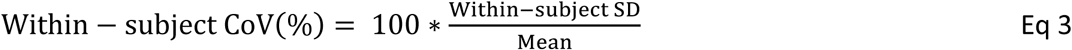

With n being the number of subjects (=14 in this study) and *x* 1 and *x* 2 are the duplicate parameter measurements for each subject.

## Results

With the total 54 examinations, data-cleaning removed 6 examinations without satisfying inclusion criteria; leading to a success rate of 89% (Fig. 2). 14 volunteers were finally included for measurement reproducibility analysis (5 males and 9 females; mean age: 25.7 years; range: 24-27 years). 68.11% of the scanned slices were finally included for the final analysis, with a mean of 5.3 slices (range: 3-10 slices) for each scan. The image series graded of ‘good quality’ presented a higher rate of acceptable fitted curve (81.2%) than slices graded ‘Fair Quality’ (62.1%).

Bland–Altman plots for PF, Dslow and Dfast each parameter are shown in Fig 5. The 95% Bland-Altman limits of agreements, mean of CoV and wCoV for repeatability and reproducibility are shown in tables 1 and 2.

**Fig 5.**
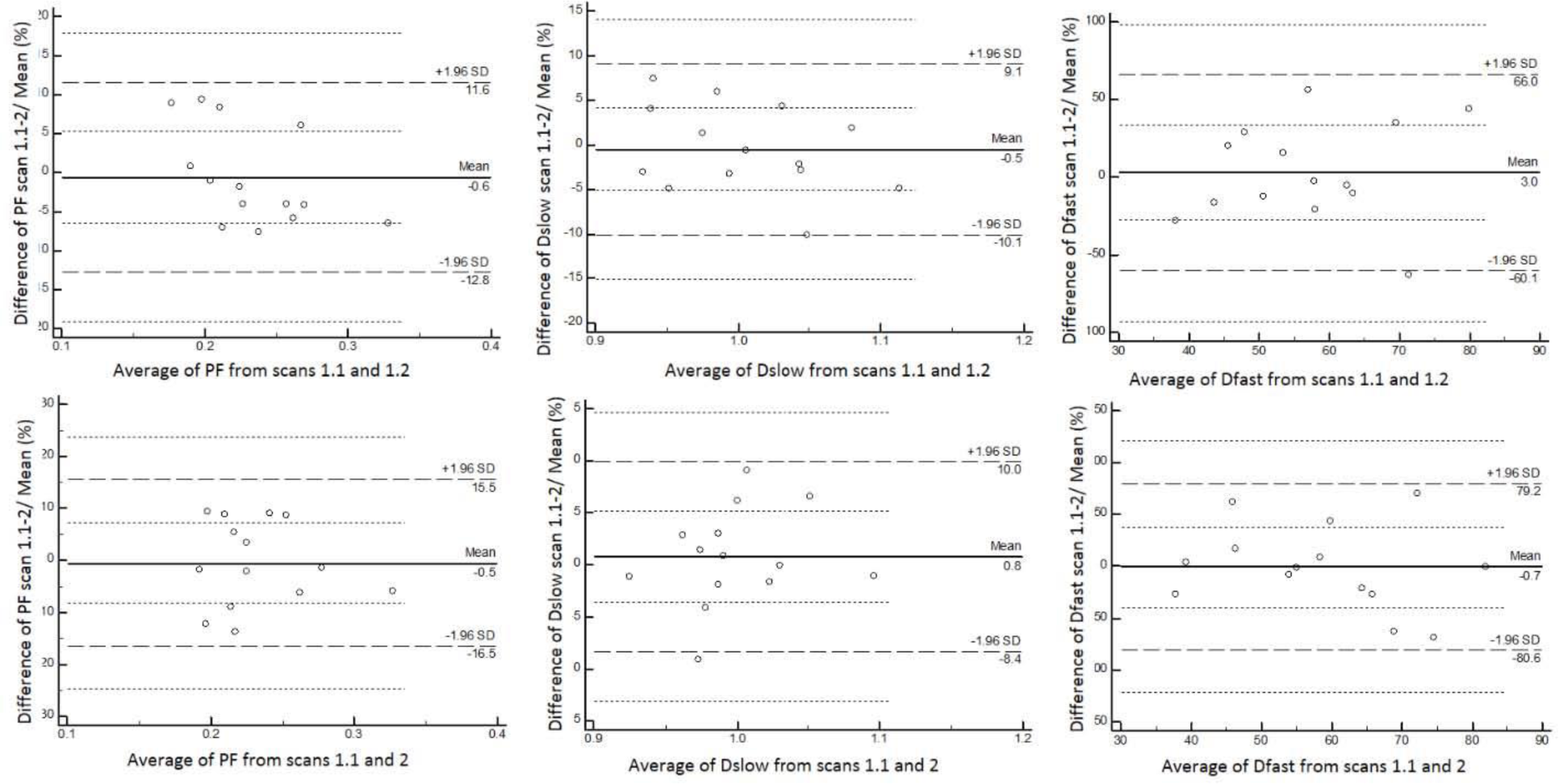
Bland–Altman plots of scan-rescan repeatability (scan 1.1 vs scan 1.2) and reproducibility (scan 1.1 vs scan 2).

**Table 1.** scan-rescan repeatability of IVIM parameters*.

| | | threshold $b=50$ | threshold $b=80$ | threshold $b=200$ |
| --- | --- | --- | --- | --- |
| **PF** | Mean CoV (% range) | 2.40 (0.13 – 7.20) | 3.81 (0.62 – 6.61) | 4.89 (0.00 – 11.49) |
|  | Within-subject CoV (%) | 3.20 | 4.24 | 6.32 |
|  | Mean Diff % (BA – LA, %) | -0.9 (-10.1 – 8.3) | -0.6 (-12.8 – 11.6) | -3.0 (-19.1 – 13.0) |
| **Dslow** | Mean CoV (% range) | 2.07 (0.30 – 5.10) | 2.86 (0.43 – 7.18) | 3.65 (0.51 – 10.34) |
|  | Within-subject CoV (%) | 2.47 | 3.36 | 4.43 |
|  | Mean Diff % (BA – LA, %) | -0.7 (-7.6 – 6.3) | -0.5 (-10.1 – 9.1) | 0.9 (-12.1 – 13.9) |
| **Dfast** | Mean CoV (% range) | 23.71 (5.01 – 49.02) | 18.16 (1.93 – 44.50) | 17.07 (0.47 – 50.39) |
|  | Within-subject CoV (%) | 28.28 | 24.88 | 26.50 |
|  | Mean Diff % (BA – LA, %) | 0.9 (-77.2 – 79.0) | 3.0 (-60.1 – 66.0) | 7.6 (-54.6 – 69.9) |

Diff: difference; BA-LA: Bland-Altman 95% limits of agreement; threshold *b*-value unit: s/mm^2^

**Table 2.** Scan-rescan reproducibility of IVIM parameters*.

| | | threshold $b=50$ | threshold $b= 80$ | threshold $b=200$ |
| --- | --- | --- | --- | --- |
| **PF** | Mean CoV (% , range) | 5.68 (0.75 – 10.94) | 4.91 (0.97 – 9.71) | 5.98 (0.60 – 13.87) |
|  | Within-subject CV (%) | 6.59 | 5.38 | 6.94 |
|  | Mean Diff % (BA – LA, %) | -0.7 (-19.9 – 18.5) | -0.5 (-16.5 – 15.5) | -1.8 (-22.2 – 18.6) |
| **Dslow** | Mean CoV (% , range) | 2.99 (0.61 – 6.75) | 2.48 (0.05 – 6.45) | 3.10 (0.58 – 6.00) |
|  | Within-subject CV (%) | 3.72 | 3.24 | 3.53 |
|  | Mean Diff % (BA – LA, %) | 0.8 (-9.7 – 11.3) | 0.8 (-8.4 – 10.0) | 1.3 (-8.4 – 11.0) |
| **Dfast** | Mean CoV (% , range) | 21.60 (1.15 – 67.31) | 21.18 (0.23 – 49.87) | 19.33 (2.31 – 72.36) |
|  | Within-subject CV (%) | 34.33 | 30.89 | 35.93 |
|  | Mean Diff % (BA – LA, %) | 3.2 (-84.0 – 90.3) | -0.7 (-80.6 – 79.2) | -4.3 (-80.1 – 71.5) |

Diff: difference; BA-LA: Bland-Altman 95% limits of agreement; threshold *b*-value unit: s/mm^2^

## Discussion

In this study, we demonstrated that IVIM parameters can have good reproducibility when evidential motion contaminated and/or poorly fitted image data are removed. Quite a few papers have addressed IVIM parameter reproducibility (10-18, 20-27). Our results broadly represent the best results ever reported.

In addition to the ‘data cleaning’ process taken in this study, a few other steps may have additionally contributed to the good results in this study. The participants were trained to avoid irregular breathing or sudden deep breathing during the examination. Sixteen *b*-values were used, which is at the up end of *b*-value number compared with published results on IVIM reproducibility (table 6). We were able to use a mean of 5.3 slices for reproducibility calculation, which is more than the number of slices used in most of the published papers on reproducibility (10-18, 20-27). The signal measurement was ROI-based method, and the IVIM parameters were calculated based on the mean signal intensity of the whole ROI. ROI-based approach allows assessing the plots of signal measurements and fitted curves for each slices, while this is not possible for pixel-based method when IVIM parameters are generated on parametric maps.

Our study has some limitations. The volunteer population in this study included only young healthy subjects. While our results may be applicable to diffused liver diseases such as hepatic fibrosis, how our approach can be applicable to focal liver lesions will require additional studies. Secondly, we didn’t ask volunteers to fast before examination, while the hepatic flow may vary depending on the fasted/prandial status (28). The reproducibility of IVIM parameters could therefore may be better when the subjects are scanned in fasted status. The data cleaning criteria presented in this study remains subjective, not precisely defined and was not automatized. An objective assessment method, including machine-based recognizing anatomical landmarks and estimation of quantification of scattering of the MRI signal intensity vs *b*-value relationship, are being explored in our laboratory to automatically and consistently assess the data acquisition quality. Another limitation is that we only tested segment-unconstrained analysis of IVIM data, while segment-unconstrained analysis remains till now the most popular approach for IVIM analysis (7), it has been suggested that Bayesian probability may perform better in fitting consistency (29, 30). Finally, in this study, six out of 54 scanned could not be used for analysis, leading to a success rate of 89%. A better sequence design allowing over-sampling of the focused liver parenchyma regions is expected to minimize the failure rate, and potentially because of the increased number of ‘sufficient quality slices’ available for averaging, will further increase the measure reproducibility.

In conclusion, we demonstrated the proof-of-principle that the scan-rescan reproducibility of IVIM parameters can potentially be good. This understanding is important for further developing IVIM technique for diagnostic clinical application.

## Acknowledgement

We thank Miss Yao Tina Li, former research student at the Chinese University of Hong Kong, for programming the image processing tool used in this study; Dr Jean-Pierre Cercueil at Department of Vascular and Interventional Radiology, François-Mitterrand Teaching Hospital, University of Bourgogne/Franche-Comté, Dijon, France, for discussions during the course of data analysis. Dr Olivier Chevallier was supported by a grant provided by the Société Française de Radiologie (SFR) together with the Collège des Enseignants de Radiologie de France (CERF).

